# Characterising open chromatin identifies novel cis-regulatory elements important for paraxial mesoderm formation and axis extension

**DOI:** 10.1101/2020.01.20.912337

**Authors:** Gi Fay Mok, Leighton Folkes, Shannon Weldon, Eirini Maniou, Victor Martinez-Heredia, Alice Godden, Ruth Williams, Grant N. Wheeler, Simon Moxon, Andrea E. Münsterberg

## Abstract

The development of multicellular organisms is exquisitely regulated through differential gene activity, which governs cell differentiation programs. However, many details of spatiotemporal control of gene regulation are still poorly understood. We used the accessibility of chick embryos to examine genome-wide signatures characterizing the progressive differentiation of paraxial mesoderm along the head-to-tail axis. Paraxial mesoderm becomes organized into repetitive units, termed somites, the hallmark of the segmented vertebrate body plan. New somite pairs form periodically as the axis extends at the posterior end. This process generates a developmental gradient within a single embryo, with anterior somites more advanced in their differentiation compared to posterior somites. Following somite formation, cell rearrangements generate compartments, comprising lineages of the musculoskeletal system, including cartilage of the vertebral column and ribs, and skeletal muscle cells of the trunk and limbs. To examine how paraxial mesoderm becomes regionalized and patterned to eventually generate these discrete lineages, we investigated dynamic changes of the transcriptome and of chromatin accessibility using RNA-seq and ATAC-seq across a spatiotemporal series along the embryonic axis. Footprint analysis uncovers differential coverage of binding sites for a number of key transcription factors known to be involved in axial patterning and differentiation, including HOX genes. Furthermore, associating accessible chromatin with nearby expressed genes identifies candidate <u>c</u>is-<u>r</u>egulatory <u>e</u>lements (CRE). As exemplars we use TCF15 and MEOX1, which are crucial for somite formation and differentiation, to experimentally validate CREs *in vivo* using fluorescent reporters. Time-lapse microscopy reveals CRE spatiotemporal activity and mutation analysis uncovers necessary upstream regulators. The CRE for MEOX1 is conserved and recognized in Xenopus. In addition, a human element is active in chicken. *In vivo* epigenome editing of TCF15 and MEOX1 CREs disrupts gene expression regulation and recapitulates phenotypic abnormalities of anterior-posterior axis extension.

## INTRODUCTION

The partitioning of paraxial mesoderm into repetitive segments, termed somites, is a key feature of vertebrate embryos. During amniote gastrulation, mesoderm cells emerge from the primitive streak and migrate in characteristic trajectories to generate axial, paraxial and lateral plate mesoderm (Iimura et al., 2007; Yang et al., 2002). Paraxial mesoderm is located on either side of the midline tissues, neural tube and notochord. As the body axis extends, it consecutively generates pairs of somites (Benazeraf and Pourquie, 2013) – epithelial spheres comprised of multipotent progenitor cells. In response to extrinsic signals, epithelial somites undergo dramatic morphogenetic changes and reorganize (Brent and Tabin, 2002; Christ et al., 2007; Gros et al., 2009; McColl et al., 2018). On the ventral side cells undergo an epithelial to mesenchymal transition (EMT) to form the sclerotome, while on the dorsal side the cells in the dermomyotome remain epithelial. From the dermomyotome edges cells transition to form the myotome, in-between the sclerotome and dermomyotome (Gros et al., 2004). Concomitantly with somite morphogenesis, the differentiation potential of somite cells becomes more restricted, with cells eventually becoming specified towards the lineages of the musculoskeletal system, including chondrocytes and skeletal muscle cells (Brent and Tabin, 2002). Overall the process of somitogenesis generates a spatiotemporal gradient of differentiation within the paraxial mesoderm along the embryonic body axis (Benazeraf and Pourquie, 2013).

In addition, somite derivatives exhibit regional differences depending on their anterior-posterior axial position. Regional identity is already established at gastrula stages and is controlled by the step-wise transcriptional activation of HOX gene expression (Kmita and Duboule, 2003; Neijts and Deschamps, 2017; Noordermeer et al., 2014). For example, members of the HOXB cluster are first activated in a temporal colinear fashion in prospective paraxial mesoderm, prior to ingression through the primitive streak (Iimura and Pourquie, 2006). The colinear activation of HOX genes culminates in nested expression domains within the paraxial mesoderm, thereby conferring regional identity along the axis (Aires et al., 2018; Iimura et al., 2009). To determine the structural features associated with colinear expression the 3D organization of HOX clusters has been investigated (Noordermeer et al., 2014). It has also been shown that posterior Wnt signalling and CDX transcription factors are important regulators of the “trunk” HOX genes in the centre of HOX clusters (Tabaries et al., 2005). In particular, CDX2 is essential for axial elongation with mutations leading to posterior truncations associated with changes in HOX expression domains (Chawengsaksophak et al., 2004). CDX activity is associated with histone acetylation and mediates chromatin accessibility of regulatory elements (Neijts et al., 2017).

Superimposed onto regional differences is the control of cell identity and differentiation and several well-characterized transcription factors serve as markers for musculoskeletal lineages. Chondrogenic cells express PAX1, PAX9 and SOX9 and dermomyotomal myogenic progenitors are characterized by PAX3 and PAX7. Committed myoblasts express MYF5 and MYOD, while MYOG and KLHL31 are markers for differentiated myocytes (Abou-Elhamd et al., 2015; Berti et al., 2015; Mok et al., 2015). Other transcriptional regulators that are important in paraxial mesoderm include TCF15 (Paraxis), a bHLH transcription factor required for somite epithelialization (Burgess et al., 1996)CDX (Caudal), which is necessary for axis elongation (Young et al., 2009); and MEOX1, which is involved in somite morphogenesis, patterning and differentiation, particularly of sclerotome derived structures (Mankoo et al., 2003; Skuntz et al., 2009). In human, mutations of MEOX1 are found in patients with Klippel-Feil Syndrome, which is associated with fusion and numerical defects in the cervical spine as well as scoliosis (Bayrakli et al., 2013; Mohamed et al., 2013).

Whilst the sequence of marker gene expression in paraxial mesoderm is well defined (Berti et al., 2015; Mok et al., 2015), the epigenetic and genomic mechanisms that control these transcriptional programs remain largely unknown. The identification of enhancers has improved through high-throughput sequencing assays and comparative genomic analysis, however, experimental validation of enhancer activity remains challenging. Here we assay spatiotemporal changes in both gene expression signatures and accessible chromatin that occur in differentiating paraxial mesoderm along the anterior-posterior axis. We define differentially accessible chromatin regions within HOX genes that are associated with regional identities. Footprint analysis reveals differential occupancy and coverage of binding sites along the axis for several transcription factors, including HOXA10, HOXA11, CDX2, LEF1 and RARA. By correlating accessible chromatin with nearby expressed genes, we identify novel cis-regulatory elements. We focus here on enhancers located upstream of TCF15 and MEOX1 and validate these *in vivo*, using electroporation of fluorescent enhancer reporters into gastrula stage chick embryos. Time-lapse imaging uncovers the onset of enhancer activation in paraxial mesoderm and mutation of candidate transcription factor motifs leads to loss of gene expression and phenotypic changes. Altogether our data provide a comprehensive resource and represent the first characterization of the accessible chromatin and gene expression landscapes in paraxial mesoderm, at different stages of patterning and differentiation of the developing musculoskeletal system.

## RESULTS

### Transcriptional profiling of developing paraxial mesoderm

To conduct genome-wide transcriptome analysis during the spatiotemporal transition of paraxial mesoderm, we collected presomitic mesoderm (PSM), epithelial somites (ES), maturing somites (MS) and differentiated somites (DS) from a Hamburger-Hamilton stage 14 (HH14) (Hamburger and Hamilton, 1951) chick embryo in triplicate. At this stage, the four most posterior somites are still epithelial, but in maturing somites cells in the ventral part undergo EMT, the dorsal dermomyotome lip forms in the epaxial domain adjacent to the neural tube and myogenic cells begin to transition into the early myotome. Differentiating somites are compartmentalized, with a primary myotome beneath the dermomyotome and a sclerotome ventrally (Christ et al., 2007; Kalcheim and Ben-Yair, 2005).

After harvesting, tissues were processed for RNA-sequencing (RNA-seq) (Figure 1A). Principal-component analysis (PCA) showed that PSM, ES, MS and DS samples cluster into three distinct groups, with MS and DS samples clustering together (Figure S1A). Differential gene expression analysis comparing PSM and ES revealed up-regulation of 713 genes and down-regulation of 583 genes; comparing ES and MS revealed up-regulation of 145 genes and down-regulation of 155 genes; and comparing MS and DS revealed up-regulation of 53 genes and down-regulation of 26 genes (Figure 1B). Additional comparisons between samples confirmed that the greatest differential was observed between PSM and somite samples, followed by the number of differentially expressed genes between the most recently formed epithelial somites and the most differentiated somites (ES vs DS)(Figure S1B).

**Figure 1.**
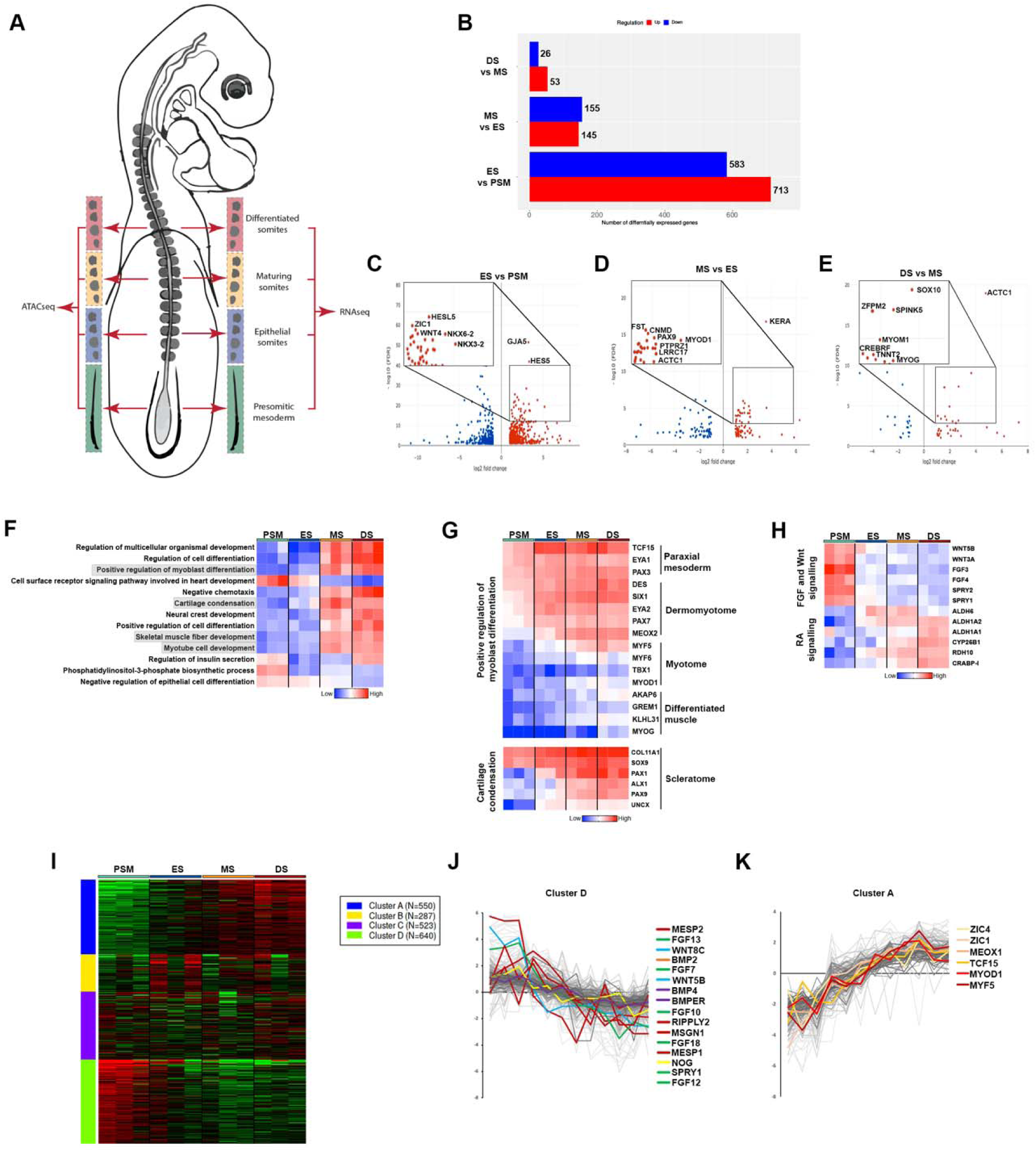
Transcriptional profiling of developing somites. (A) Schematic representation of HH14 chick embryo with presomitic mesoderm (PSM), epithelial somite (ES), maturing somite (MS) and differentiated somite (DS) dissected for RNA-seq and ATAC-seq. (B) Bar chart showing number of differentially expressed (upregulated, red; downregulated, blue) genes comparing PSM with ES, ES with MS and MS and DS. (C-E) Volcano plots showing enriched genes (Log Fold Change >1.5) comparing PSM with ES, ES with MS and MS and DS. (F) Heatmap showing GO terms associated with PSM or DS enriched genes. (G) Clusters of highly correlated genes identified for myoblast differentiation, cartilage condensation, (H) Wnt, FGF and retinoic acid (RA) signalling pathways are shown in heatmap. (I) Heatmap showing k-means linear enrichment clustering across PSM, ES, MS and DS. (J) Cluster D and (K) Cluster A are shown with genes labelled.

Previously described somite transcription factors (TF), such as NKX6-2, NKX3-2, ZIC1 and HES5, were highly enriched in ES compared to PSM, as well as the gap junction protein GJA5 (Connexin 40). Marker genes important for myogenic (MyoD1 and ACTC1) and chondrogenic (PAX9, FST) cell lineages were enriched in MS compared to ES. Novel markers for chondrocytes (Chondromodulin, CNMD), bone homeostasis (Leucine-rich repeat containing, LRRC17) and cartilage, (Keratan sulfate proteoglycan Keratocan, KERA) were also identified (Figure 1D). Myogenin (MYOG), a TF involved in differentiation of muscle fibres was enriched in DS compared to MS, as was expression of the neural crest cell (NCC) transcription factor SOX10, likely due to NCCs migrating in close proximity to and through the rostral half of differentiating somites. Other genes highly expressed in DS include the serine protease inhibitor, SPINK5; Troponin T2 (TNNT2) and Myomesin (MYOM1), encoding important proteins of the sarcomere and mediating contraction; CREBRF, a negative regulator of the endoplasmic reticulum stress response and ZFPM2, a zinc finger TF (Figure 1E).

The functional clustering by gene ontology (GO) terms of differentially expressed genes across all four stages reveals enrichment of biological processes involved in cell differentiation in DS and MS versus PSM and ES samples. In particular, genes involved in myoblast differentiation, cartilage condensation, skeletal muscle fibre development and myotube cell development were upregulated (Figure 1F). Further analysis of genes involved in positive regulation of myoblast differentiation shows that they display dynamic expression across the four groups and include classic markers for different stages of paraxial mesoderm differentiation. Genes differentially expressed in somite compartments include in the dermomyotome and myotome: MYF5, MYF6 and MYOD1, whilst MYOG and KLHL31 are associated with differentiated muscle. Classic markers for chondrogenesis and cartilage condensation within the sclerotome include the transcription factors, SOX9, PAX1 and PAX9, and the extracellular matrix component, COL11A1 (Figure 1G). Functional clustering of differentially expressed genes also reveals enrichment of signalling pathways involved in anterior-posterior pattern formation. These pathways are expressed in an opposing fashion and include the FGF and Wnt signalling pathways, which are highly expressed in PSM and the retinoic acid (RA) signalling pathway, which is more highly expressed in somite samples (Figure 1H).

We next used weighted gene co-expression network analysis (WGCNA) (Langfelder and Horvath, 2008; Langfelder et al., 2008) to characterise gene co-expression clusters across the four samples of the top 400 differentially expressed genes (Figure 1I). We identified 4 clusters and discovered genes that were decreasing (Cluster D) (Figure 1J) or increasing (Cluster A) (Figure 1K) in expression across the spatiotemporal domains, from PSM to DS. Genes in Cluster D feature components of FGF (FGF13, FGF7, FGF10, FGF18, SPRY1), BMP (BMP2, BMP4, BMPER, NOG) and WNT (WNT8C, WNT5B) signalling pathways in addition to classic PSM markers such as MESP2, RIPPLY2, MSGN1 and MESP1, which are known to be important for somitogenesis. Genes in Cluster A feature markers of cellular differentiation programs such as ZIC1, ZIC4, MEOX1, TCF15 and include the myogenic regulatory factors, MYOD1 and MYF5.

### Profiling chromatin accessibility dynamics in paraxial mesoderm along the anterior-posterior axis

Next, we wanted to identify genomic regulatory elements that control paraxial mesoderm and somite differentiation programs. To do this, we used ATAC-seq (Assay for Transposase-Accessible Chromatin using sequencing) (Buenrostro et al., 2013) to map chromatin accessibility across the paraxial mesoderm along the axis, in PSM, ES, MS and DS (Figure 1A). Distinct chromatin accessibility profiles were evident at different stages of somite development, indicative of the dynamic progression of axial development. Principle component analysis (PCA) showed a high reproducibility between biological triplicates of each sample type (Figure S2G-L), but dynamic changes in chromatin accessibility were observed between them. Using DiffBind (Ross-Innes et al., 2012; Stark and Brown, 2011), we show the densities and clustering of the differentially accessible chromatin regions (peak-sites) (rows), as well as the sample clustering (columns) for PSM against ES, ES against MS and MS against DS. We identified differentially accessible peaks with differential densities showing clusters of peak sites with distinct patterns of chromatin accessibility levels for PSM against ES (Figure 2A), ES against MS (Figure 2B) and MS against DS (Figure 2C). MA plots show the highest number of differentially accessible peaks is evident when comparing PSM and ES (n=27692, Figure 2D). The number of differentially accessible peaks is lower when comparing ES against MS and MS against DS (n=4670, n=1965, Figure 2E and F). This is in line with the transcriptome data, where greater differences were seen between PSM and ES compared to the differences observed between different stages of somite maturation.

**Figure 2.**
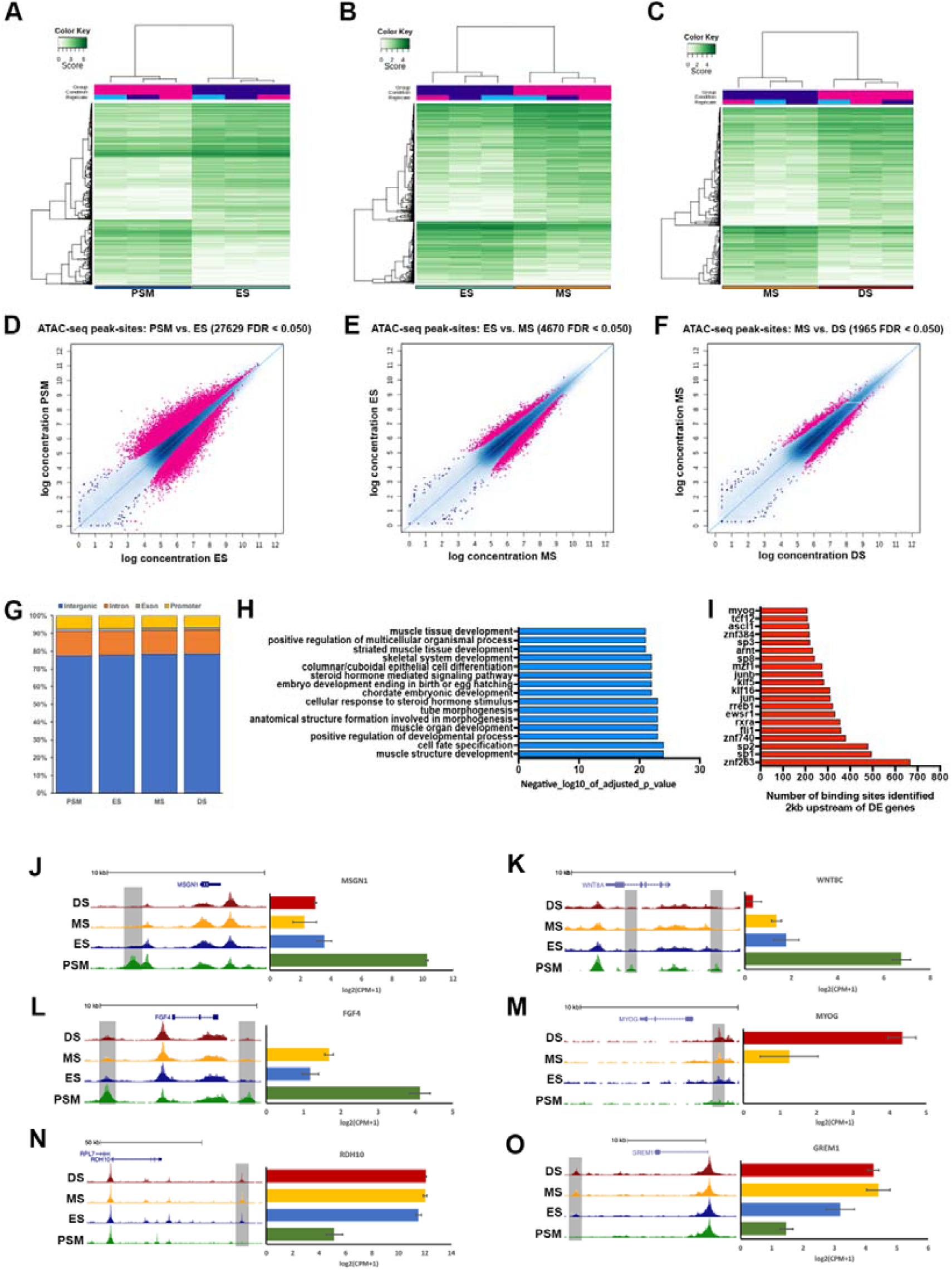
Genome-wide profile of chromatin accessibility dynamics during somite development. (A) Correlation heat maps of accessible chromatin regions (ATAC-seq peak-sites) comparing PSM and ES, (B) ES and MS and (C) MS and DS. (D) MA plots of significantly differential peak-sites (pink) in PSM with ES, (E) ES with MS and (F) MS with DS. (G) Bar plot showing proportions of total genome sequence of peaks in PSM, ES, MS and DS. Most peaks lie in intergenic and intron regions. (H) GO terms associated with enriched transcription factors in DS compared to PSM. (I) Number of transcription factor binding sites identified 2kb upstream of differentially expressed genes in DS compared to PSM. (J-O) Examples of ATAC-seq accessible regions that define PSM (Msgn1, Wnt8c, Fgf4) or DS (MyoG) or developing somites (Rdh10, Grem1). Gene expression from mRNA-seq (error bars = SEM) is shown in the bar charts on the right.

The genomic distribution of accessible regions was similar in all four sample types, the majority, 78%, were located in intergenic regions, approximately 12% were located in introns, 2% in exons and 8% of accessible regions were associated with promoter regions (Figure 2G). Functional terms associated with predicted transcription factor (TF) binding sites that were enriched in accessible peaks in DS compared to PSM included cell fate specification and terms related to morphogenesis or skeletal myogenesis (Figure 2H). Consistent with the latter, we identified >200 binding sites for myogenin (MYOG) that are located within accessible chromatin peaks within 2 kb of genes differentially expressed in DS, where skeletal muscle differentiation occurs (Figure 2I). The MYOG motif is well conserved across mouse and human, thus is likely to be conserved across avian species also. Other TFs enriched identified include bHLH proteins (TCF12, ASCL1, ARNT1), of which TCF12 is expressed in skeletal muscle, is part of the canonical Wnt pathway and implicated as a transcriptional repressor in colorectal cancer (Lee et al., 2012). Specificity proteins, Sp1, Sp2, Sp3 and Sp8, are zinc finger proteins known to interact with bHLH proteins such as MyoD (Biesiada et al., 1999). Sp1 and Sp3 bind to GC and GT boxes and can be displaced from these sequences by KLF16, a Krüppel-like zinc finger protein for which binding sites are also enriched. Other zinc finger TFs include ZNF384, ZNF740 and ZNF263, which are involved in the regulation of cell differentiation genes including those relevant to musculoskeletal development. For example ZNF384 regulates extracellular matrix genes MMP1, MMP3, MMP7 and COL1A1 (Nakamoto et al., 2000); ZNF740 recruits the chromatin regulator HDAC1 to the SMAD4-DNA complex and prevents the recruitment of the transcriptional activators CREBBP and EP300 (Yang et al., 2015) and ZNF263 is involved in adipogenesis (Ambele and Pepper, 2017). The Ewing sarcoma RNA binding protein 1 (EWSR1) has an N-terminal transcriptional activation domain and a C-terminal RNA-binding domain. EWSR1 regulates gene expression, cell signaling, RNA processing and transport. Chimeric proteins resulting from chromosomal translocations between EWSR1 and various transcription factor genes, including the genes encoding ZNF384 and FLI1 (Delattre et al., 1992; Grunewald et al., 2015), are involved in tumorigenesis, such as acute leukemia or Ewing sarcoma in bones and bone connective tissues. Furthermore, the binding motif for retinoic acid receptor alpha (RXRA) is enriched. RXRA is a nuclear receptor, which can act as a transcriptional repressor or transcriptional activator, depending on the context.

Next, we identified differentially accessible peaks that were open specifically in PSM or in somite samples, ES, MS or DS. We hypothesise that these could represent putative enhancers. For example, differentially accessible peaks identified flanking genes highly expressed in the PSM included a peak downstream of MSGN1 present in PSM and not in ES, MS or DS (Figure 2J); a peak downstream of WNT8C and a peak within intron 1 present in PSM but not in somite tissues (Figure 2K); and peaks upstream and downstream of FGF4 present in PSM and low or absent in other tissues (Figure 2L). For the muscle differentiation gene, MYOG, a peak was identified upstream of the gene in DS, MS and interestingly also in ES, but not in PSM (Figure 2M). For RDH10, which is associated with RA signalling and highly expressed in somites but less abundant in PSM, a differential peak is identified in ES, MS and DS and not in PSM (Figure 2N). Similarly, for GREM1, an antagonist of BMP signalling highly expressed in developing somites, a differential peak is identified downstream of the gene that is present in ES, MS and DS but not in PSM. In most cases, chromatin accessibility correlates well with gene expression and in some cases, it precedes the detection of transcripts, e.g. MYOG. However, putative enhancer activities of these differential peaks remain to be confirmed experimentally.

### Identification of differential footprints during somite development

To further interrogate the accessible chromatin landscape during somite development, we used HINT-ATAC (Li et al., 2019) to discover differential transcription factor footprints in regions of open chromatin identified in PSM, ES, MS or DS. Initially we focussed on PAX3, an important TF that regulates the myogenic program and is highly expressed during somite development (Figure 3A). HINT-ATAC revealed an increase in the number of PAX3 footprints in ES open chromatin when compared to PSM (Figure 3B). The number of PAX3 footprints increased further in MS and DS when compared to PSM (Figure 3C, D) suggesting that there is a greater coverage of bound sites in maturing and differentiating somites and consistent with the role of PAX3 in myogenic progenitors in the dermomyotome. Another key somite TF, TWIST2 (also known as DERMO1), is important for epithelial-to-mesenchymal transition (EMT) during somitogenesis. TWIST2 is highly expressed in the paraxial mesoderm and increases as somites differentiate (Figure 2E). The number of genome wide TWIST2 footprints are very similar in PSM and ES (Figure 2F), however, the number of footprints increases in MS and DS when compared to PSM (Figures 2G, H).

**Figure 3.**
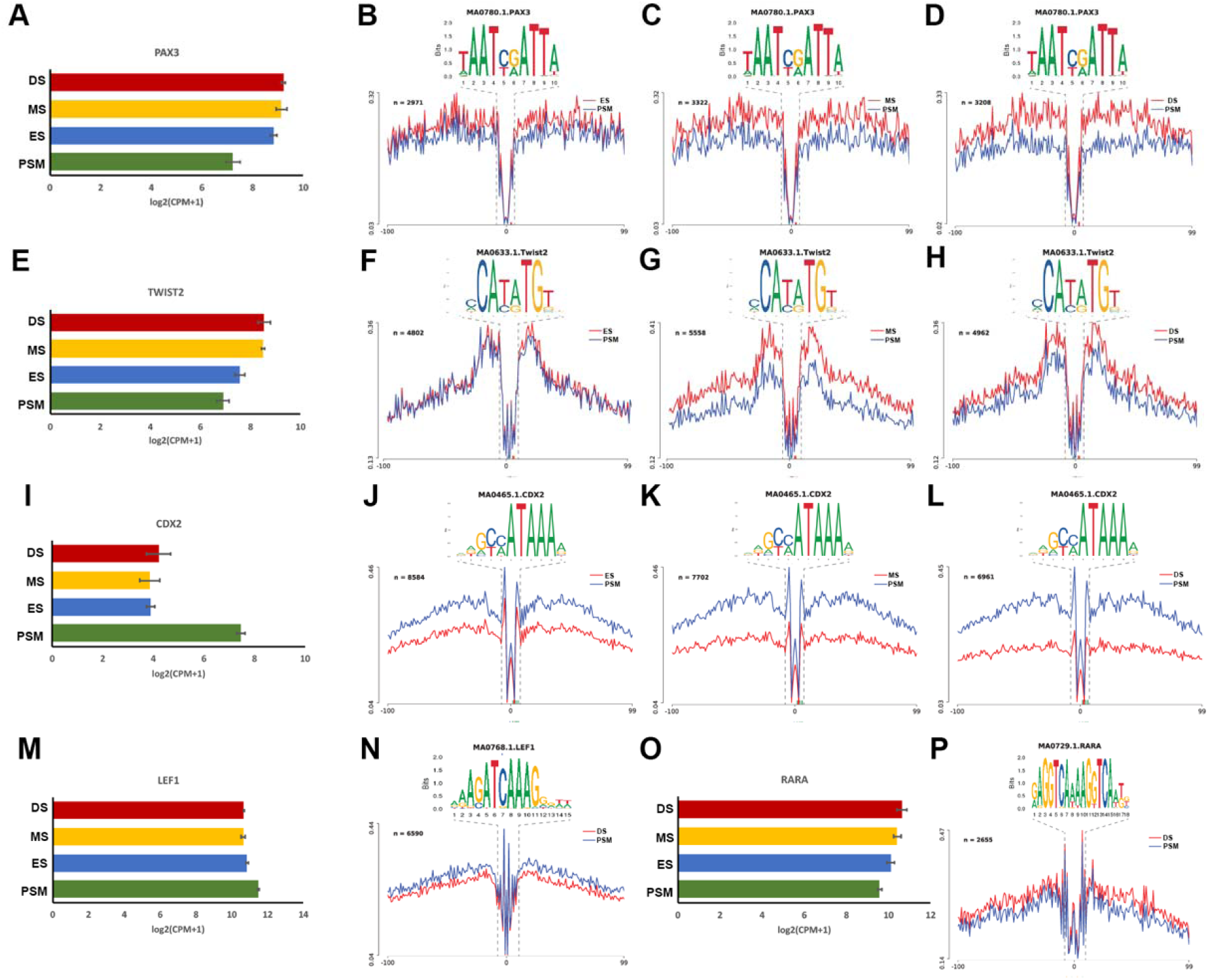
Differential footprints identified during somite development. (A) Gene expression from mRNA-seq (error bares = SEM) for PAX3. (B) Differential footprinting for Pax3 motif comparing PSM and ES, (C) PSM and MS and (D) PSM and DS. (E) Gene expression for TWIST2. (F) Differential footprinting for TWIST2 motif comparing PSM and ES, (G) PSM and MS and (H) PSM and DS. (I) Gene expression for CDX2. (J) Differential footprinting for CDX2 motif comparing PSM and ES, (K) PSM and MS and (L) PSM and DS. (M) Gene expression and (N) differential footprinting for LEF1 comparing PSM and DS. (O) Gene expression and (P) differential footprinting for RARA comparing PSM and DS.

The CDX2 transcription factor is a readout for posterior WNT signalling and has been implicated in defining neuro-mesodermal progenitors (Metzis et al., 2018). In addition, CDX2 is essential for axial elongation (Young et al., 2009) and is highly expressed in the PSM (Figure 3I). Consistent with high levels of WNT signalling activity in the PSM, HINT-ATAC identified a greater number of CDX2 footprints in open chromatin in this region when compared to ES, MS and DS (Figure 3J-L). Similarly, LEF1, a transcriptional effector for canonical WNT signalling, is highly expressed in the PSM (Figure 3M), although it is also expressed in developing somites, where it becomes restricted to the myotome (Schmidt et al., 2004; Schmidt et al., 2000). We identified a greater number of LEF1 footprints in the PSM when compared to DS, consistent with the more restricted expression of LEF1 in the latter (Figure 3N).

Next, we investigated footprints for retinoic acid receptor alpha, RARA, which mediates RA (retinoic acid) signalling. Consistent with its high expression in somites (Figure 3O), HINT-ATAC identified fewer RARA footprints in PSM compared to DS (Figure 3P). Overall, the coverage patterns observed for CDX2, LEF1 and RARA are consistent with the opposing expression patterns for WNT and RA pathways observed within the paraxial mesoderm along the anterior-posterior axis (Figure 1H).

### Chromatin accessibility and differential TF footprints in the HoxA cluster

We next examined the HOXA cluster, which imposes regional identity along the anterior-posterior axis via the colinear expression of its members, and determined how HOXA gene expression patterns correlate with the accessible chromatin landscape. RNA sequencing determined expression levels of each member of the HOXA cluster in PSM, ES, MS and DS (Figure 4A). Their expression reflects the organisation of the genes within the cluster: the more 3’ located genes have a more anterior expression boundary compared to the genes located more 5’, which are restricted more posteriorly. Accordingly, we find that HOXA1, HOXA2, HOXA3, HOXA4, HOXA5 and HOXA6 are all highly expressed across the length of the axis: in PSM, ES, MS and DS. A small decrease in HOXA7 gene expression is detected in DS, with more pronounced decreases observed for HOXA9, HOXA10, HOXA11 and HOXA13, which are also reduced progressively in MS and ES. The colinear pattern of gene expression correlates with differentially accessible chromatin within the HOXA cluster (Figure 4B). Accessible chromatin regions are seen in PSM, ES, MS and DS near the promoter of HOXA1, HOXA2, HOXA3, HOXA4, HOXA5 and HOXA6. However accessible chromatin for HOXA7 is reduced at the promoter in DS compared to PSM, ES and MS. For the more posteriorly restricted genes, HOXA9, HOXA10, HOXA11 and HOXA13 accessible chromatin peaks are reduced in ES, MS and DS compared to PSM, which correlates with their reduced expression. To investigate the impact of the dynamic changes in HOXA gene expression along the anterior-posterior axis, we next explored the number of TF footprints for HOXA2, HOXA5, HOXA10 and HOX11 in PSM and DS (Figure 3C-F). We observed the same number of footprints for HOXA2 and HOX5 when comparing PSM and DS, however, a significant decrease in coverage is detected for HOXA10 and HOXA11 footprints in anterior DS compared to PSM. This reveals a strong association between gene expression levels along the anterior-posterior axis and the genome-wide coverage of HOXA binding sites.

**Figure 4.**
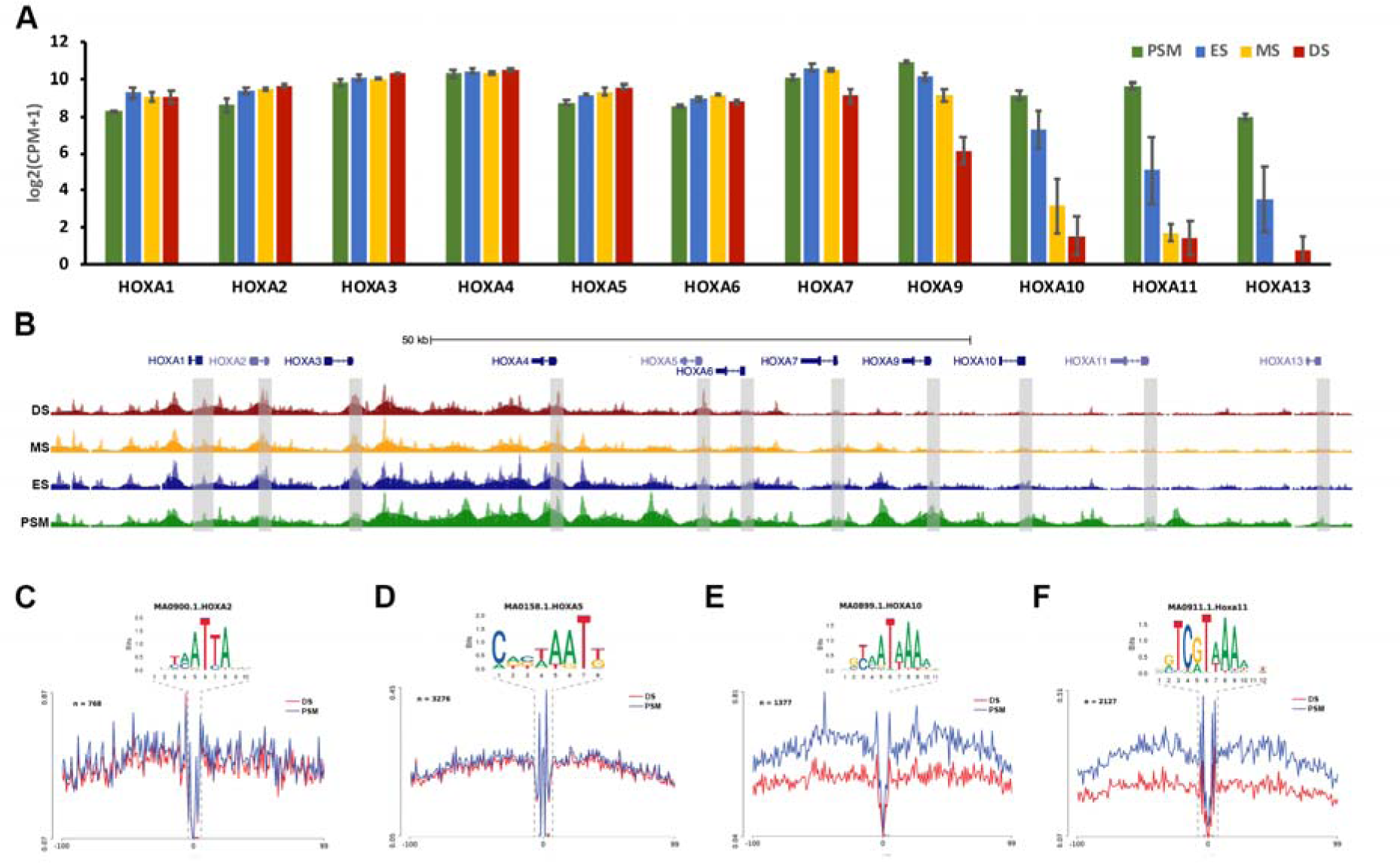
Chromatin accessibility and differential footprints for HoxA cluster. (A) Gene expression from mRNA-seq for HoxA cluster (error bars = SEM). (B) Genome browser views of ATAC-seq profile at HoxA cluster. PSM ATAC is shown in green, ES shown in blue, MS shown in yellow and DS in red. Grey boxes indicate promoter region of each HoxA gene. Differential footprinting for (C) HoxA2, (D) HoxA5, (E) HoxA10 and (F) HoxA11 between PSM and DS.

### Identification and validation of paraxial mesoderm-specific regulatory elements

To validate and further characterize potential enhancers active specifically in somites we carried out embryo electroporation (Chapman et al., 2001). We focus here on TCF15 and the homeodomain transcription factor, MEOX1, two classic markers for paraxial mesoderm specification and somite patterning. To identify enhancers for these genes, we examined open chromatin peaks flanking the gene within 10kb. Identified peaks representing candidate *cis*-regulatory elements (CRE) were cloned upstream of the herpes simplex virus thymidine kinase (HSV-TK) minimal promoter, driving expression of a stable Citrine reporter (Williams et al., 2019). Electroporation targeted the prospective mesoderm of gastrula-stage HH3+ embryos (Figure S3A), and reporter gene expression profiles were monitored until HH11.

We identified two CREs upstream of TCF15 (Figure 5A), which showed spatially restricted enhancer activities. For the first element, TCF15 Enh-1 (1500 bp) we observed activity in the PSM, in all somites and in the notochord (Figure 5B). The second element, TCF15 Enh-2 (700 bp), showed activity mainly in PSM and somites, as well as some activity in lateral plate mesoderm (LPM) (Figure 5C and D). *In situ* hybridization shows that expression of TCF15 is restricted to PSM and somites (Figure 5F) (Bothe and Dietrich, 2006). Therefore, it is not clear at present why TCF15 Enh-1 and TCF15 Enh-2 drive reporter expression also in the notochord and LPM. It is possible that repressive elements that limit enhancer activity are missing, or that the enhancers are driving a different gene in these cells. We think it is less likely that accumulation of Citrine reveals sites of gene expression that cannot be detected by *in situ* hybridization. Time-lapse movies for TCF15 Enh-2 show Citrine fluorescence is first detected in a HH6 embryo in prospective paraxial mesoderm cells as they converge towards the midline (Movie S1). Strong signal is seen in the first somite at HH7 and subsequently in all newly formed somites, as well as the PSM and prospective paraxial mesoderm cells.

**Figure 5.**
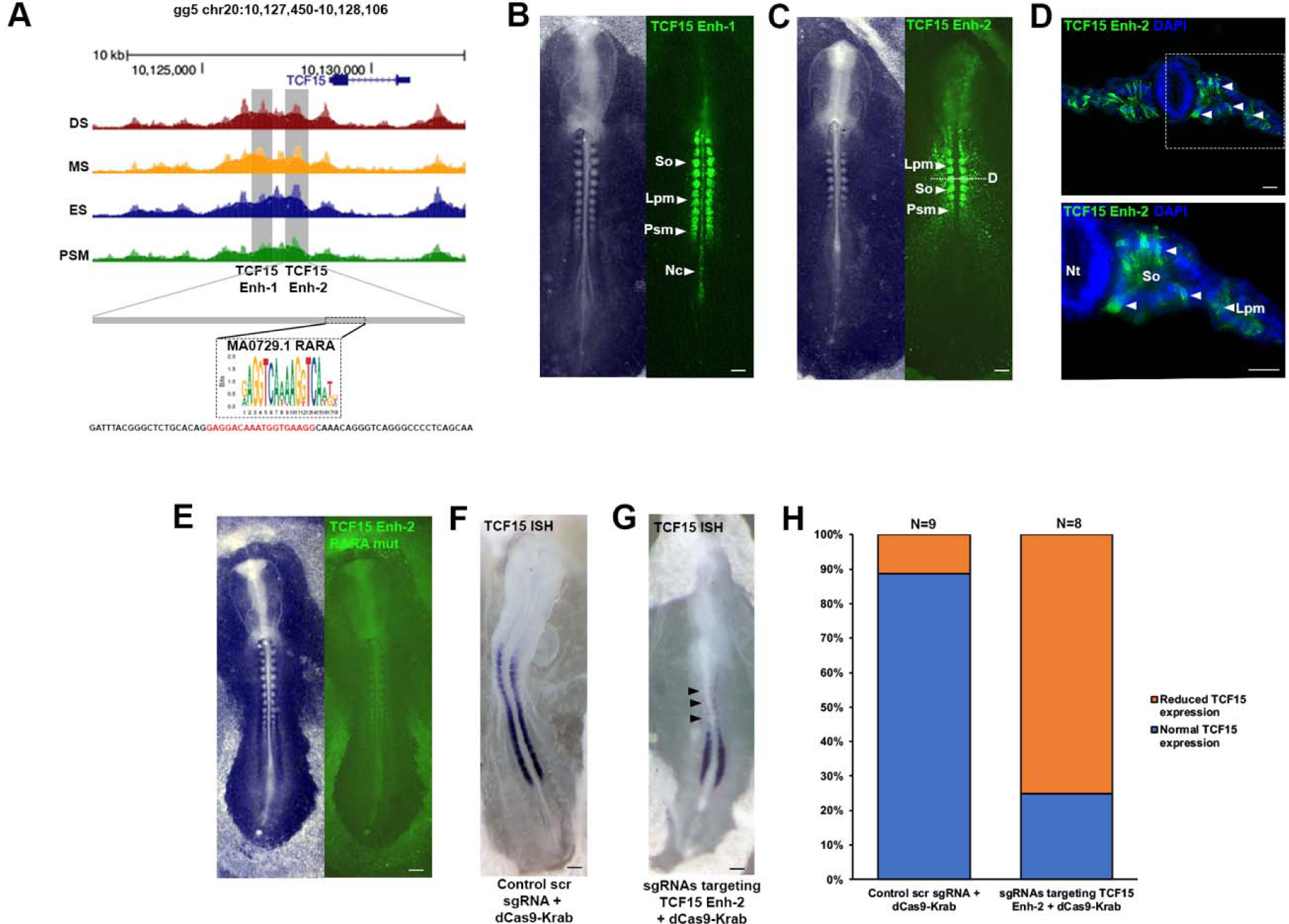
Identification of paraxial mesoderm-specific enhancers. (A) ATAC-seq profile at Tcf15 locus. Grey boxes indicate putative enhancers identified (TCF15 Enh-1 and TCF15 Enh-2). RARA footprint identified within Tcf15 Enh-2. Mutant reporter sequence for TCF15 Enh-2 RARA mutant. (B) Tcf15 Enh-1 and (C) Tcf15 Enh-2 reporter expression in presomitic mesoderm (PSM), notochord (Nc), somites (So) and lateral plate mesoderm (Lpm). (D) Transverse sections of Tcf15 Enh-2 immunostained for Citrine showing expression in somites and lateral plate mesoderm, white dashed line in (C) indicates location of section. Nuclei stained with DAPI (blue). (E) Tcf15 Enh-2 Citrine reporter with RARA binding site mutation displays lack of expression in somites. (F, G) Epigenome engineering using dCas9-Krab with (F) control scrambled sgRNAs and (G) sgRNAs targeting endogenous RARA binding site followed by wholemount *in situ* hybridisation for TCF15. (H) Percentage of embryos with normal (blue) or reduced (orange) TCF15 *in situ* expression after electroporation of control scrambled sgRNA with dCas9-Krab or sgRNAs targeting TCF15 Enh-2 RARA binding site with dCas9-Krab.

Because reporter activity observed with TCF15 Enh-2 reflects more closely the spatial gene expression pattern of TCF15, we next sought to identify TFs that regulate this element. HINT-ATAC identifies a TF footprint for retinoic acid receptor alpha (RARA) within TCF15 Enh-2, consistent with RARA expression and coverage of binding sites across the anterior-posterior axis (Figure 3O, P). Introducing mutations into the RARA binding site (Figure 5A) leads to loss of reporter activity in the embryo (Figure 5E), suggesting RARA is indeed required to activate TCF15 Enh-2. To determine the potential significance of RARA-mediated regulation of TCF15 Enh-2 *in vivo*, we used the conventional dCas9-KRAB repressor to modify the endogenous enhancer (Williams et al., 2018). Two CRISPR guide RNAs (gRNA) designed to target the repressor to the TCF15 Enh-2 RARA binding site, or scrambled gRNA controls were electroporated together with dCas9-KRAB (Figure 5F, G). Detection of TCF15 expression by *in situ* shows that epigenomic modification of the endogenous TCF15 Enh-2 alone leads to reduced TCF15 expression and concomitantly a drastic truncation of the body axis (N=6/8 embryos, Figure 5G), whilst control scrambled gRNAs/dCas9-KRAB repressor has no effect on TCF15 expression or axis elongation (N=8/9 embryos, Figure 5F, H). These data suggest that retinoic acid (RA) signalling is crucial for TCF15 gene expression as RARA binding-site perturbation results in disruption of anterior-posterior axis elongation. This is consistent with mouse mutants of TCF15 or mutants affecting RA signalling, in which epithelial somite formation is disrupted and the embryonic axis is truncated (Burgess et al., 1996; Ghyselinck and Duester, 2019; Vermot and Pourquie, 2005). It has been shown that Wnt signalling is important for TCF15 expression in early somites (Linker et al., 2005), however, there is no evidence of direct regulatory interactions. Although we cannot exclude the possibility that Wnt signalling, via LEF/b-catenin, contributes to TCF15 expression and that this is mediated via a different CRE, which alone is not sufficient. It is also worth noting that the conservation of TCF15 Enh-1 and Enh-2 across mammalian species is poor (data not shown).

MEOX1 is an important TF for early somite patterning and differentiation (Mankoo et al., 2003; Skuntz et al., 2009). We examined accessible regions of chromatin that were evolutionary conserved at sequence level between chicken, Zebra finch, American alligator, Chinese softshell turtle, lizard, human and mouse. This approach identified one candidate CRE of 1095 kb, approximately 1kb upstream of MEOX1 (Figure 6A). This element displayed enhancer activity, with expression of the Citrine reporter restricted to the PSM and all somites (Figure 6B, C). Time-lapse movies reveal Citrine fluorescence, which is first detected in the prospective paraxial mesoderm cells of a HH6 embryo. At HH7 signal is detected in the first somite and subsequently in all newly formed somites, as well as the PSM and prospective paraxial mesoderm cells. Overall the pattern is consistent with MEOX1 gene expression detected by *in situ* (Figure 6D, F) (Movie S2). We identified two predicted TF footprints within the enhancer, one for FOXO1 and one for ZIC3 (Figure 6A). We next determined their requirement for the activation of fluorescent reporter expression. When individual sites for FOXO1 or ZIC3 were mutated, reporter activity is still observed (data not shown). However, mutation of both sites leads to loss of reporter activity (Figure 6E). This suggests both TFs are able to activate this CRE and either FOXO1 or ZIC3 alone is sufficient. To investigate the significance of this element, we modified the endogenous enhancer using four guide RNAs (gRNAs) to target the dCas9-KRAB repressor to the MEOX1 Enh. Scrambled gRNAs with dCas9-KRAB were used as control (Figure 6F-H). Using a probe to detect MEOX1 transcripts *in situ* shows that MEOX1 Enh enhancer perturbation leads to loss of gene expression (Figure 6G), suggesting this element is required. As the MEOX1 Enh is highly conserved amongst amniote taxa - birds, reptiles and mammals (Figure 6I), we next asked whether the homologous mammalian sequences are active in chick. We found that a human MEOX1 Enh, isolated from HeLa cells, is able to drive fluorescent reporter gene expression in somites. Activity is also detected in lateral plate and presegemented mesoderm (Figure 6J). The human MEOX1 Enh sequence included conserved FOXO1 and ZIC3 binding sites, and mutation of both sites leads to loss of reporter activity (Figure 6K) suggesting these sites are required for enhancer activity. Therefore, we propose transcriptional regulation of the MEOX1 enhancer is highly conserved between human and chick. Interestingly, the MEOX1 Enh sequence was not found in fish or amphibians (Figure 6I). However, when we injected the chick MEOX1 Enh reporter into one cell of *Xenopus laevis* embryos at the 2-cell stage, we observed Citrine fluorescence in paraxial mesoderm in early neurula stages (NF stage 14), during early mesoderm formation (NF stage 25) and in somite-derived muscle fibres (NF stage 33 and stage 42) (Figure 6L). The MEOX1 Enh with mutations in the FOXO1 and ZIC3 binding sites shows no fluorescent activity (data not shown). This suggests that the MEOX1 Enh can be activated by the same regulatory mechanism in amphibians, most likely via FOXO1 and ZIC3, even though the CRE is not conserved in the same location in the *Xenopus laevis* genome.

**Figure 6.**
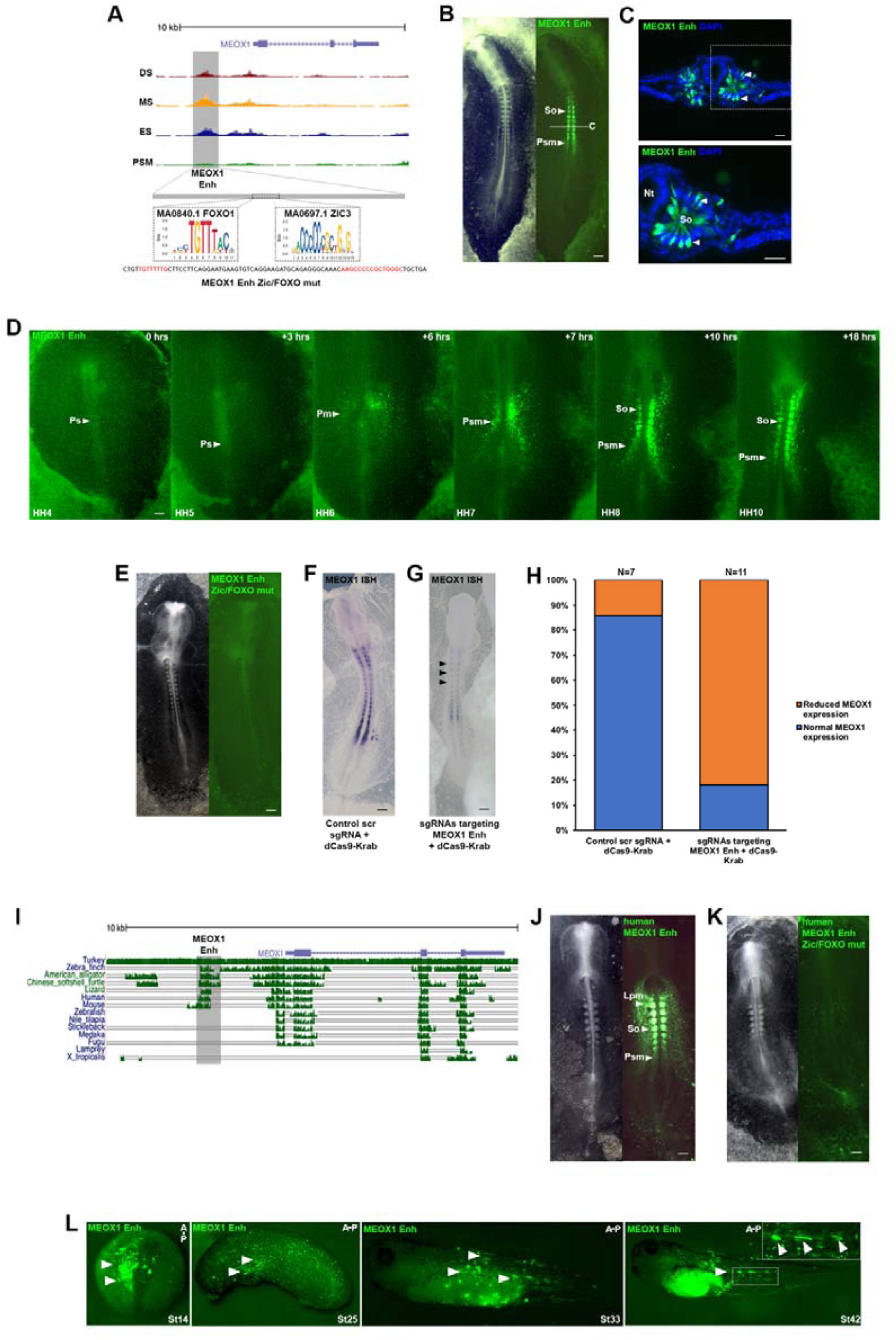
Identification of a conserved regulatory element for Meox1. (A) ATAC-seq profile at MEOX1 locus. Grey box indicates putative enhancer identified. FOXO1 and ZIC3 footprints identified within enhancer element. Mutant reporter sequence for MEOX1 Enh ZIC/FOXO mutant. (B) MEOX1 Enh reporter expression identified in presomitic mesoderm (PSM) and in somites (So) along anterior-posterior axis. (C) Transverse sections of MEOX1 Enh immunostained for Citrine showing expression in epithelial somites, white dashed line indicates location of section. Nuclei stained with DAPI (blue). (D) Still photographs from a time-lapse movie of MEOX1 Enh with the primitive streak (Ps) indicated. Fluorescent activity first observed in paraxial mesoderm (PSM) at HH6 and continuous expression in the paraxial mesoderm (PSM) at HH7 prior to expression in somites (So) at HH8 and HH10. (E) FOXO1 and ZIC3 binding sites mutated in MEOX1 Enh Citrine reporter which displayed lack of expression in somites and presomitic mesoderm. Epigenome engineering using dCas9-Krab with (F) control scrambled sgRNAs and (G) sgRNAs targeting endogenous FOXO1 and ZIC3 binding sites followed by wholemount *in situ* hybridisation for MEOX1. (H) Percentage of embryos with normal (blue) or reduced (orange) MEOX1 expression after injection and electroporation of control scrambled sgRNA with dCas9-Krab or sgRNAs targeting MEOX1 Enh FOXO1 and ZIC3 binding sites with dCas9+Krab. (I) Genomic alignment of chick MEOX1 locus. Top indicates the scales in kilobase. Blue thin vertical lines below gene names indicate the protein coding exons. Clustered green vertical lines indicate the presence of sequence similarity with the sequences of different species, indicated on the left at each position. The height of bars indicates extent of conservation in different species: turkey, zebrafinch, American alligator, Chinese softshell turtle, Lizard, Human, Mouse, Zebrafish, Nile tilapia, Stickleback, Medaka, Fugu, Lamprey and Xenopus tropicalis. Grey box indicates MEOX1 Enh. (J) Human MEOX1 Enh reporter expression evident in presomitic mesoderm and somites. (K) FOXO1 and ZIC3 binding site mutations in human MEOX1 Enh Citrine reporter results in lack of expression. (L) Chick MEOX1 Enh injected into *Xenopus laevis* embryos at 2-cell stage shows Citrine reporter activity in paraxial mesoderm (St14) and in early somites and muscle fibres (St25, St33, St42).

## DISCUSSION

Extension of the vertebrate body axis is driven by the addition of new segments at the posterior end of the embryo. During gastrulation, presumptive paraxial mesoderm cells ingress through the primitive streak (Iimura and Pourquie, 2006; Yang et al., 2002). In addition, paraxial mesoderm formation depends on a bi-potential population of neuro-mesodermal progenitor cells (NMP). In response to high levels of Wnt3a and CDX family members, these progenitors commit to mesoderm fates and give rise to neck, trunk and tail structures (Aires et al., 2018).

Here we provide molecular profiles of paraxial mesoderm of cranial and trunk regions by RNA-seq and ATAC-seq (Buenrostro et al., 2013). This identifies differential gene expression consistent with axial patterning and differentiation, including the appearance of chondrogenic and myogenic markers in more anterior differentiating somite samples (Figure 1D, E, G, K) (Bothe and Dietrich, 2006; Christ et al., 2007; Gros et al., 2004; Kalcheim and Ben-Yair, 2005; Mok et al., 2015). Components of signalling pathways involved in anterior posterior axis patterning are also differentially expressed, such as FGF, Wnt and RA pathways (Figure 1H, J) (Dubrulle and Pourquie, 2004; Vermot and Pourquie, 2005). Furthermore, we uncover genome-wide dynamic changes in chromatin accessibility across the spatiotemporal series. Using HINT-ATAC, an improved method to predict transcription factor binding sites (TFBS) with footprintes (Li et al., 2019), we reveal for the first time differential coverage along the anterior-posterior axis for several binding sites, including sites for RARA and LEF1, transcriptional effectors of the RA and Wnt pathways (Figure 3M-P). The observed coverage patterns correlate well with gene expression and with the known functions of RA and Wnt signalling in anterior-posterior axis patterning.

We also observe differential chromatin accessibility along the anterior-posterior axis in the HOXA cluster (Figure 4B), as well as differential expression (Figure 4A) and footprints (Figure 4C-F) of HOXA family members. These patterns are consistent with the role of HOX clusters in the regionalization of axial structures (Neijts and Deschamps, 2017; Noordermeer et al., 2014). In addition CDX2, which is essential for posterior axis elongation (Chawengsaksophak et al., 2004), is highly expressed in PSM and has greater coverage of footprints in PSM compared to somite samples (Figure 3I-L). PSM samples correspond to thoracic axial levels and this region is defined by central HOX genes, which are regulated by CDX proteins (Neijts et al., 2017; Tabaries et al., 2005).

Predicting enhancer gene interactions remains challenging, although new computational methods are becoming available. For example, the recent activity-by-contact model indicates that very long-range interactions are rare (Fulco et al., 2019; Gaffney, 2019). Here we selected accessible chromatin regions within 10 kb of the transcription start site. This approach combined with experimental validation and time-lapse imaging in gastrula stage chick embryos identified novel *cis*-regulatory elements for TCF15 and MEOX1, both of which are in close proximity of the transcription start site (Figures 5 and 6). Footprints identified candidate TF binding sites and dCas9-Krab epigenome modification (Williams et al., 2018) leads to loss of expression suggesting these CREs are essential. In addition, *in vivo* epigenome modification of TCF15-Enh2 and the MEOX1 CREcaused axial elongation phenotypes (Figure 5G, H, Figure 6G, H), which are consistent with mouse mutants (Burgess et al., 1996; Mankoo et al., 2003) and human Klippel-Feil patients (Bayrakli et al., 2013; Mohamed et al., 2013).

Taken together we provide a comprehensive data set across a series of paraxial mesoderm samples, which has uncovered novel CREs important for vertebrate anterior-posterior axis formation and will enable further interrogation and data-mining.

## MATERIALS AND METHODS

### Chicken embryos

Fertilised chicken eggs (Henry Stewart & Co.) were incubated at 37°C with humidity. Embryos were staged according to Hamburger and Hamilton (Hamburger and Hamilton, 1951). All experiments were performed on chicken embryos younger than 14 days of development and therefore were not regulated by the Animal Scientific Procedures Act 1986.

### Embryo dissection

Hamburger and Hamilton stage 14 (HH14) embryos were dissected into Ringers solution in silicon lined petri dishes and pinned down using the extra-embryonic membranes. Ringers solution was replaced with Dispase (1.5mg/ml) in DMEM 10mM HEPES pH7.5 at 37°C for 7 minutes prior to treatment with Trypsin (0.05%) at 37°C for 7 minutes. The reaction was stopped with Ringers solution with 0.25% BSA. The PSM, ES, MS and DS were carefully dissected away from neural and lateral mesoderm tissue using sharp tungsten needles.

### RNA extraction, library preparation and sequencing

For ES, MS and DS, consecutive four somites were dissected. Tissues were placed into RLT lysis buffer. RNA was extracted using Qiagen RNAeasy kit (Cat no. 74104) and DNase treated (Qiagen Cat no 79254) for removal of DNA. Libraries were prepared and sequenced on the Illumina HiSeq4000 platform (75bp paired end) at the Earlham Institute. A minimum of three biological replicates for each stage were used for analysis.

### ATAC, library preparation and sequencing

PSM, ES, MS and DS samples were dissected as stated above. Cell dissociation was performed using a protocol adapted from (Williams et al., 2019). Briefly, tissues were dissociated with Dispase at 37°C for 15 minutes with intermittent pipetting to attain a single cell suspension with 0.05% Trypsin at 37°C for a final 5 minutes at 37°C. The reaction was stopped, and cells were re-suspended in Hanks buffer (1X HBSS, 0.25% BSA, 10mM HEPES pH8). Cells were centrifuged at 500 g for 5 minutes at 4°C, re-suspended in cold Hanks buffer, passed through 40μm cell strainers (Fisher Cat no. 11587522), and further centrifuged at 500 g for 5 minutes at 4°C. Pelleted cells were re-suspended in 50ul Hanks buffer, kept on ice and processed for ATAC library preparation. ATAC was performed using a protocol adapted from (Buenrostro et al., 2013; Williams et al., 2019). Briefly, cells were lysed in cold lysis buffer (10mM Tris-HCl, Ph7.4, 10mM NaCl, 3mM MgCl2, 0.1% Igepal) and tagmentation performed using Illumina Nextera DNA kit (FC-121-1030) for 30 minutes at 37°C on a shaking thermomixer. Tagmented DNA was purified using Qiagen MinElute kit (Cat no. 28004) and amplified using NEB Next High-Fidelity 2X PCR Mast Mix (Cat no. M0543S) for 11 cycles as follows: 72°C, 5 minutes; 98°C, 30 seconds; 98°C, 10 seconds; 63°C, 30 seconds; 72°C, 1 minute. Library preparation was complete after further clean up using Qiagen PCR MinElute kit (Cat no. 28004) and Beckman Coulter XP AMPpure beads (A63880). Tagmentation size was assessed using Agilent 2100 Bioanalyser. Libraries were quantified with Qubit 2.0 (Life Technologies) and sequenced using paired-end 150bp reads on the Illumina HiSeq4000 platform at Novogene UK. A minimum of three biological replicates for each stage were used for analysis.

### Enhancer cloning

Chick genomic DNA (gRNA) was extracted from HH14 embryos using Invitrogen Purelink gDNA extraction kit (Cat no. K1820-00). Human genomic DNA was isolated from HeLa cells. Putative enhancers were amplified using primers with specific sequence tails (see Supplemental Table 1) to enable cloning into reporter vector using a modified GoldenGate protocol (Engler et al., 2009) under the following conditions: 94°C, 3 minutes; 10 cycles of 94°C, 15 seconds; 55°C, 15 seconds; 68°C, 3 minutes, 25 cycles of 94°C, 15 seconds; 63°C, 15 seconds; 68°C, 3 minutes; and final step of 72°C, 4 minutes. Amplicons were purified using Qiagen PCR Cleanup (Cat no. 28104) and pooled with pTK nanotag reporter vector with T4 DNA ligase (Promega) and BsmBI (NEB) restriction enzyme. This reaction was prepared for T4-mediated ligation and BsmBI digestion under the following conditions: 25 cycles of 37°C, 2 minutes; 16°C, 5 minutes; a single step of 55°C, 5 minutes; and a final step of 80°C, 5 minutes. For mutagenesis of specific sites in enhancers we utilised FastCloning methodology (Li et al., 2011). Primer sequences are detailed in Supplemental Table 1.

### CRISPR-mediated enhancer repression

sgRNAs specific for MEOX1 Enh and TCF15 Enh-2, or a scrambled control were cloned into a chicken pU6-3 vector as previously described (Williams et al., 2018). For enhancer repression, sgRNAs and dCas9-Krab were electroporated *ex ovo* (Williams et al., 2018). Primer sequences are detailed in Supplemental Table 1.

### Embryo preparation and *ex ovo* electroporation

Hamburger and Hamilton (HH3+) embryos were captured using the filter paper based easy-culture method. Briefly, eggs were incubated for approximately 20 hours, a window was created using forceps, the embryo and yolk were transferred into a dish and thin albumin above and around the embryo was removed using tissue paper. A circular filter paper ring was placed on top, excised and transferred into a separate dish containing Ringers solution and excess yolk was removed. The embryo was then transferred into a dish containing albumin-agar and ready for electroporation with the ventral side up (Chapman et al., 2001). Plasmid DNA was injected between the membrane and embryo to cover the whole epiblast, electroporated used 5 pulses of 5V, 50ms on, 100ms off. Thin albumin was used to seal the lids of dishes and embryos were cultured at 37°C with humidity to the desired stage.

### Cryosectioning and immunostaining

Embryos were fixed in 4% PFA (paraformaldehyde) for 2 hours at room temperature (RT) or at 4°C overnight, washed 3x 10 minutes in PBS. Embryos were transferred into 30% sucrose/PBS overnight at 4°C prior to 3x 10 minutes washes in OCT before final embedding of OCT in dry ice. Cryosectioning was performed at 15 μm thickness. Sections were washed in 3x 15 minutes PBS and 1x 15 minutes in PBS/0.5% Triton X-100 prior to blocking in 5% goat serum and 5% BSA in PBS for 1 hour at RT. Incubation with primary antibody for rabbit anti-GFP (1:200, Torrey Pines Biolabs Cat no. TP401) at 4°C overnight, followed by 3x 10 minute washes in PBS and incubation with secondary antibody AlexaFluor-568-conjugated donkey anti-rabbit IgG (1:500, ThermoFisher Cat no. A21206) for 1 hour at RT. Sections were washed 3x 10 minute in PBS and 1x wash with PBS and DAPI (Sigma-Aldrich) at 0.1mg/ml in PBS.

### Wholemount *in situ* hybridisation

Whole-mount *in situ* hybridization using antisense RNA probes for *Meox1* (a gift from Baljinder Mankoo, King’s College London UK) and TCF15 (a gift from Susanne Dietrich, University of Portsmouth UK) was carried out as described previously (Goljanek-Whysall et al., 2011).

### Live imaging of enhancer reporter

Embryos cultured in six-well cell culture plates (Falcon) were time-lapse-imaged on an inverted wide-field microscope (Axiovert; Zeiss). Brightfield and fluorescent images were captured every 6 min for 20–24 h, using Axiovision software as described in (Song et al., 2014). At the end of the incubation, most embryos had reached stage HH10-11.

### Image analysis

Sections were visualized on an Axioscope with Axiovision software (Zeiss). Whole mount embryos were photographed on a Zeiss SV11 dissecting microscope with a Micropublisher 3.5 camera and acquisition software or Leica MZ16F using Leica Firecam software. Live imaging datasets were analysed in FIJI/ImageJ.

### ATAC-seq Processing

Adaptors were removed from raw paired-end sequencing reads and trimmed for quality using Trim Galore! (v.0.5.0) (Krueger, 2015) a wrapper tool around Cutadapt (Martin, 2011) and FastQC (Andrews, 2010). Default parameters were used. Quality control (QC) was performed before and after read trimming using FastQC (v.0.11.6) (Andrews, 2010) and no issues were highlighted from the QC process. Subsequent read alignment and post-alignment filtering was performed in concordance with the ENCODE project’s “ATAC-seq Data Standards and Prototype Processing Pipeline” for replicated data (https://www.encodeproject.org/atac-seq/). In brief, reads were mapped to the chicken genome galGal5 assembly using bowtie2 (v.2.3.4.2) (Langmead and Salzberg, 2012). The resultant Sequence Alignment Map (SAM) files were compressed to the Binary Alignment Map (BAM) version on which SAMtools (v.1.9) (Li et al., 2009) was used to filter reads that were unmapped, mate unmapped, not primary alignment or failing platform quality checks. Reads mapped as proper pairs were retained. Multi-mapping reads were removed using the Python script assign_multimappers provided by ENCODE’s processing pipeline and duplicate reads within the BAM files were tagged using Picard MarkDuplicates (v.2.18.12) [http://broadinstitute.github.io/picard/] and then filtered using SAMtools. For each step, parameters detailed in the ENCODE pipeline were used. From the processed BAM files, coverage tracks in bigWig format were generated using deepTools bamCoverage (v 3.1.2) (Ramirez et al., 2016) and peaks were called using MACS2 (v.2.1.1) (Feng et al., 2012) (parameters -f BAMPE -g mm -B --nomodel --shift -100 --extsize 200). Coverage tracks and peaks (narrow peak format) were uploaded to the UCSC Genome Browser (Kent et al., 2002) as custom tracks for ATAC-seq data visualization.

### Differential Accessibility and Footprinting

Analysis of ATAC-seq for differential accessibility was carried out in R (v.3.5.1) (Team, 2018) using the DiffBind package (v.2.8.0) (Ross-Innes et al., 2012; Stark and Brown, 2011) with default parameter settings. Differential accessibility across samples was calculated using the negative binomial distribution model implemented in DEseq2 (v1.4.5) (Love et al., 2014). Computational footprinting analysis was conducted across samples using HINT-ATAC which is part of the Regulatory Genomic Toolbox (v.0.12.3) (Li et al., 2019) also using default parameter settings and the galGal5 genome.

### RNA-seq Differential Expression Analysis

Adaptors were removed from raw paired-end sequencing reads and trimmed for quality using Trim Galore! (v.0.5.0) using default parameters. Quality control was performed before and after read trimming using FastQC (v.0.11.6) and no data quality issues were identified after checking the resultant QC reports. Processed reads were mapped to galGal5 cDNA using kallisto (v.0.44.0) (Bray et al., 2016). Resultant quantification files were processed by custom java code to generate an expression matrix. Differential expression, GO term and pathway analyses were then conducted using DESeq2 (Love et al., 2014) and default settings within the iDEP (v.9.0) (Ge et al., 2018) web interface.GO term analysis used PGSEA method for GO Biological Process with a minimum of 15 and maximum of 2000 geneset and <0.2 FDR.

### Xenopus embryo microinjection

All experiments were carried out in accordance with relevant laws and institutional guidelines at the University of East Anglia, with full ethical review and approval, compliant to UK Home Office regulations. To obtain *X. laevis* embryos, females were primed with 100 units of PMSG and induced with 500 units of human chorionic gonadotrophin (hcG). Eggs were collected manually and fertilised *in vitro*. Embryos were de-jellied in 2% L-cysteine, incubated at 18°C and microinjected in 3% Ficoll into 1 cell at the 2 cell stage in the animal pole with 5 nL of enhancer reporter plasmid at 400 ng/μL or GFP capped RNA as control. Embryos were left to develop at 23°C. Embryo stageing is according to Nieuwkoop and Faber normal table of Xenopus development. GFP capped RNA for injections was prepared using the SP6 mMESSAGE mMACHINE kit, 5 ng was injected per embryo.

### Data availability

Raw sequencing data for this study is stored at the Sequence Read Archive (SRA) using the BioProject accession: PRJNA602335.

## ACKNOWLEDGEMENTS

We thank all members of the Münsterberg and Wheeler labs for helpful discussion. Dr Timothy Grocott for discussions, Ronce Saputil and undergraduate project students for assistance with enhancer analysis. GFM, LF, EM and VMH were supported by BBSRC (BB/N007034/1) and MRC project grants (MR/R000549/1) to AM; SW and AG were supported by studentships funded by the UKRI Biotechnology and Biological Sciences Research Council Norwich Research Park Biosciences Doctoral Training Partnership to GW and AM.

## FIGURES AND LEGENDS

**Supplemental Figure 1.**
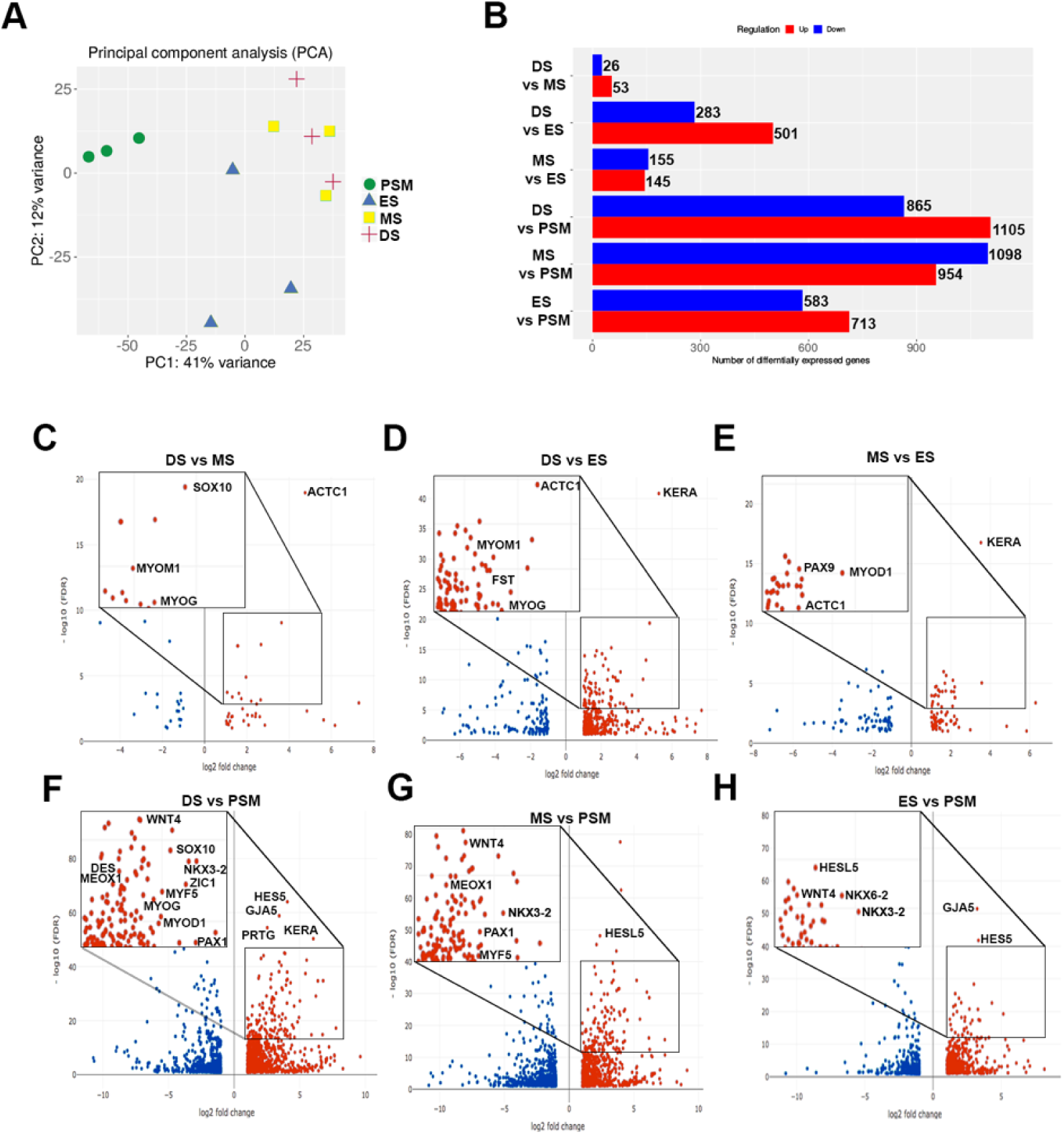
Transcriptional profiling of developing somites. (A) PCA plot of PSM, ES, MS and DS replicates. (B) Number of differentially expressed (upregulated, red; downregulated, blue) genes comparing PSM, ES, MS and DS. (C-H) Volcano plots showing enriched genes (Log Fold Change >1) comparing DS vs MS, (D) DS vs ES, (E) MS vs ES, (F) DS vs PSM, (G) MS vs PSM and (H) ES vs PSM.

**Supplemental Figure 2.**
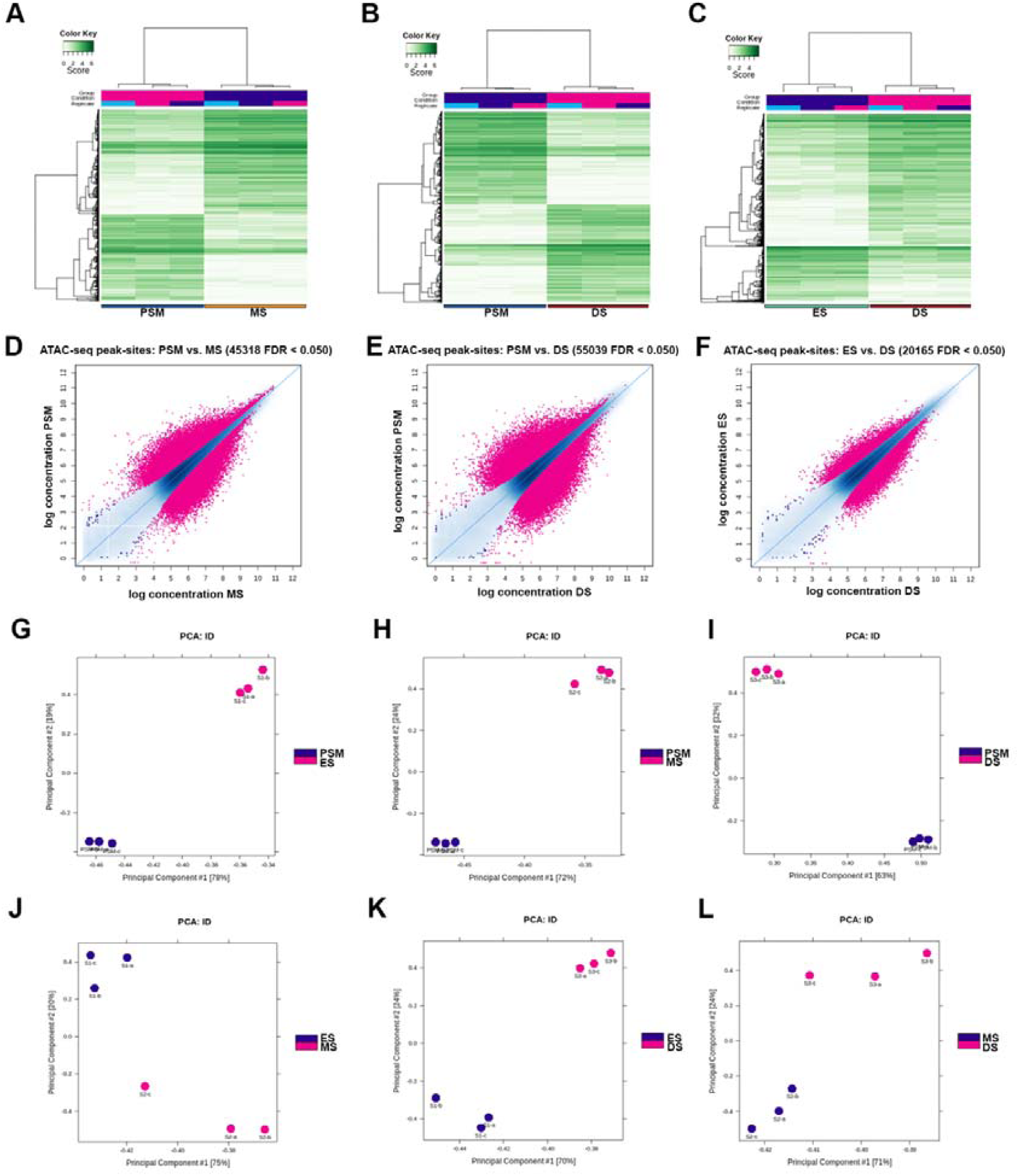
Genome-wide profile of chromatin accessibility dynamics during somite development. (A) Correlation heat maps of accessible chromatin regions (ATAC-seq peak-sites) comparing PSM and MS, (B) PSM and DS and (C) ES and DS. (D) MA plots of significantly differential peak-sites in PSM with MS, (E) ES with DS and (F) ES with DS. (G-L) PCA plots using only the differentially accessible peak-sites using an FDR threshold of 0.05.

**Supplemental Figure 3.**
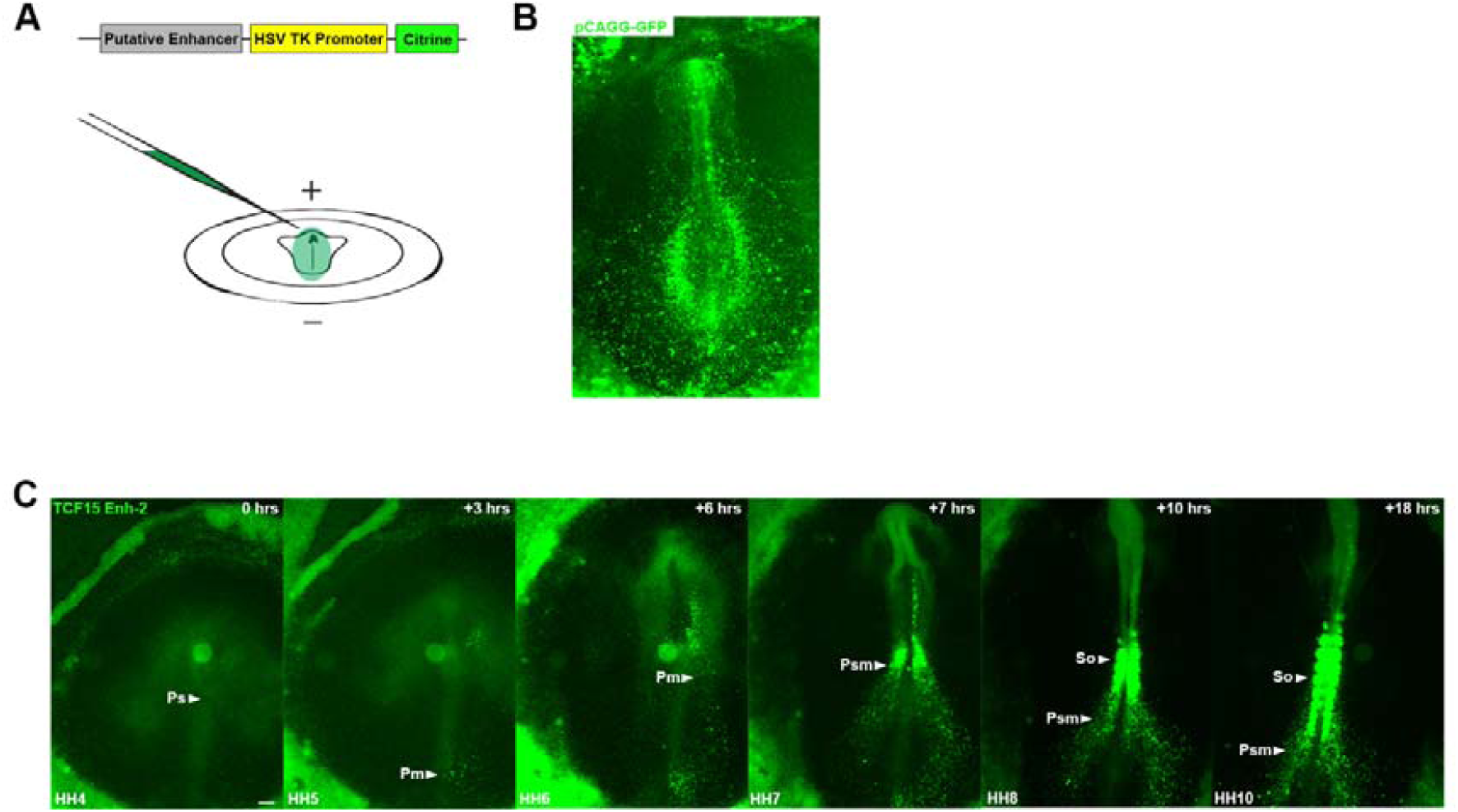
(A) Schematic representation of putative enhancer Citrine reporter construct. Injection of constructs into HH3+ chick embryos with positive electrode at the top and negative electrode at the bottom. (B) Electroporation of pCAGG-GFP at HH3+, embryo developed for 20 hours. (C) Time-lapse movie of TCF15 Enh-2 with the primitive streak (Ps) indicated. Fluorescent activity first observed in paraxial mesoderm (Pm) at HH6 and continuous expression in the paraxial mesoderm (Psm) at HH7 prior to expression in somites (So) at HH8 and HH10.

**Supplemental Table 1.**
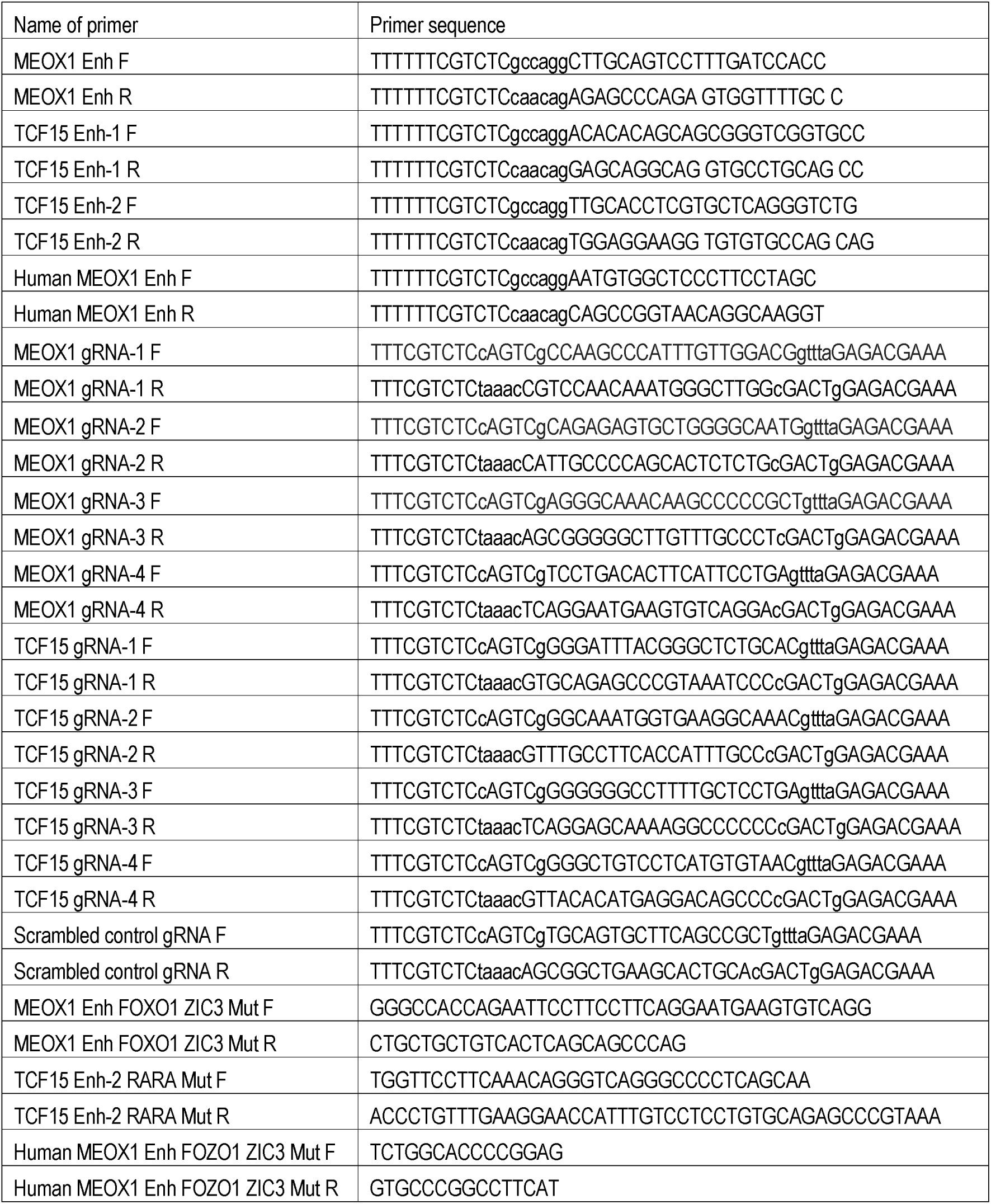

